# Rapid generation of ventral spinal cord-like astrocytes from human iPSCs for modeling non-cell autonomous mechanisms of lower motor neuron disease

**DOI:** 10.1101/2021.12.13.472474

**Authors:** Vincent Soubannier, Mathilde Chaineau, Lale Gursu, Ghazal Haghi, Anna Kristyna Franco Flores, Guy Rouleau, Thomas M Durcan, Stefano Stifani

## Abstract

Astrocytes play important roles in the function and survival of neuronal cells. Dysfunctions of astrocytes are associated with numerous disorders and diseases of the nervous system, including motor neuron diseases such as amyotrophic lateral sclerosis (ALS). Human induced pluripotent stem cell (iPSC)-based approaches are becoming increasingly important for the study of the mechanisms underlying the involvement of astrocytes in non-cell autonomous processes of motor neuron degeneration in ALS. These studies must account for the molecular and functional diversity among astrocytes in different regions of the brain and spinal cord. It is essential that the most pathologically-relevant astrocyte preparations are used when investigating non-cell autonomous mechanisms of either upper or lower motor neuron degeneration in ALS. In this context, the main aim of this study was to establish conditions enabling rapid and robust generation of physiologically-relevant ventral spinal cord-like astrocytes that would provide an enhanced experimental model for the study of lower motor neuron degeneration in ALS. Neural progenitor cells with validated caudal and ventral features were derived from human iPSCs and differentiated into astrocytes, which were then characterized by examining morphology, markers of ventral spinal cord astrocytes, spontaneous and induced calcium transients, and astrogliosis markers. Efficient and streamlined generation of human iPSC-derived astrocytes with molecular and biological properties similar to physiological astrocytes in the ventral spinal cord was achieved. These induced astrocytes express markers of mature ventral spinal cord astrocytes, exhibit spontaneous and ATP-induced calcium transients, and lack signs of overt activation. Human iPSC-derived astrocytes with ventral spinal features offer advantages over more generic astrocyte preparations for the study of both ventral spinal cord astrocyte biology and the involvement of astrocytes in mechanisms of lower motor neuron degeneration in ALS.

## Introduction

Astrocytes are a morphologically and functionally heterogenous group of glial cells in the mammalian nervous system. They perform a number of functions, such as structural and trophic support of neurons, homeostasis of the blood brain barrier, participation in synaptogenesis and plasticity, and involvement in neuroinflammatory processes.^1–3^ Astrocytes in different regions of the nervous system exhibit morphological and functional diversity reflecting their specific developmental origins and integration into different cellular microenvironments in the brain or spinal cord.^4–9^. The most frequently described example of morphological diversity among astrocytes is the presence of two major subtypes, termed protoplasmic and fibrous astrocytes, which are typically located in the gray matter and white matter, respectively.^4–9^ These morphological differences are associated with molecular and functional diversity. Specifically, protoplasmic astrocytes are in close contact with synapses and actively regulate synapse formation, maturation and function. On the other hand, white matter fibrous astrocytes are involved in mechanisms underlying axonal biology and myelination.^9–12^ These functional diversities are correlated with defined molecular traits. For instance, protoplasmic and fibrous astrocytes differ in the expression level of the intermediate filament protein, glial fibrillary acidic protein (GFAP), which is significantly higher in fibrous astrocytes than in protoplasmic astrocytes.^8–12^

The existence of multiple types of astrocytes with diverse molecular properties and functions underscores the importance of selecting the most informative experimental model systems when investigating defined biological mechanisms involving astrocytes in the developing and adult nervous system. Achieving this objective is particularly challenging when studying the roles of human astrocytes in health and disease, given the recognized difficulties in establishing primary cultures of human astrocytes. The advent of induced pluripotent stem cell (iPSC)-based approaches has addressed some of these limitations by providing efficient and reliable approaches to generate physiologically-relevant human astrocytes (e.g., references 13-17, to cite only a few). However, the diversity of astrocytes along the different axes of the nervous system poses a significant challenge to research aiming to investigate the multiple roles of astrocytes using cells derived from human iPSCs. The success of these investigations depends in large part on the ability to generate the specific astrocyte subtypes that are physiologically relevant to the biological processes under study, in both healthy and diseased conditions.

A compelling example of the importance of studying the appropriate astrocyte subtypes in the right context is provided by amyotrophic lateral sclerosis (ALS), a motor neuron disease that causes degeneration of both brain (upper) and brainstem/spinal cord (lower) motor neurons.^18–21^

Motor neuron degeneration in ALS is not only caused by cell-autonomous cell death mechanisms but can also result from non-cell autonomous processes involving non-neuronal cells, including astrocytes.^22–24^ It is believed that astrogliosis induced in response to initial insults to motor neurons plays neuroprotective roles at first but then gradually progresses towards neuroinflammatory functions that exacerbate neuronal degeneration. It is hypothesized that astrocytes exert deleterious effects on motor neurons in ALS as a consequence of either loss of supportive functions or gain of toxic functions (or a combination of both).^22–27^ Upper and lower motor neurons interact with specific subgroups of astrocytes found in either the dorsal forebrain or ventral brainstem/spinal cord, respectively. The specialized cross-talk of defined motor neurons with particular astrocyte subtypes contributes to the different functions performed by upper and lower motor neurons, and is believed to also play roles in the mechanisms of upper or lower motor neuron degeneration.^28–30^ Thus, understanding the contributions of human astrocytes to upper and lower motor neuron degeneration necessitates the implementation of experimental strategies that can generate pathophysiologically-relevant astrocyte subtypes with the appropriate rostrocaudal and dorsoventral identities.^31^ In this regard, available protocols to derive astrocytes with validated characteristics of astrocytes located in the ventral half of the spinal cord (ventral spinal astrocytes) are scarce and time-consuming.^31^

Here, we describe a protocol enabling efficient and timesaving generation from human iPSCs of astrocytes displaying molecular and biological properties of ventral spinal astrocytes. These preparations are expected to provide an advanced experimental model system for the study of spinal cord astrocyte biology and the involvement of astrocytes in mechanisms of lower motor neuron degeneration in ALS and other motor neuron diseases.

## Materials and Methods

### Human induced pluripotent stem cells

Human iPSC line AIW002-02 was established from peripheral blood mononuclear cells obtained from a 37-year-old male through retrovirus reprogramming using the CytoTune-iPS 2.0 Sendai Reprogramming kit (Thermo-Fisher Scientific; Waltham, MA, USA; Cat. No. A16518).^32^ The cell line was established at the Montreal Neurological Institute-Hospital through procedures conducted under Ethical Review Board approval by the McGill University Health Centre. Undifferentiated state of human iPSCs was routinely assessed by testing for expression of the stem cell markers NANOG and OCT4 and by quality control profiling as described previously.^32^

### Derivation of neural progenitor cells from human iPSCs

Human iPSCs at low passage number were cultured in mTeSR medium (STEMCELL Technologies; Vancouver, BC, Canada; Cat. No. 85850) in 10-cm culture dishes (Thermo-Fisher Scientific; Cat. No. 353003) coated with Matrigel (Thermo-Fisher Scientific; Cat. No. 08-774-552) until they reached 70%-80% confluence. To generate neural progenitor cells (NPCs), iPSCs were dissociated with Gentle Cell Dissociation Reagent (STEMCELL Technologies; Cat. No. 07174), followed by seeding of 2-3x10^6^ cells onto T25 flasks (Thermo-Fisher Scientific; Cat. No.

12-556-009), coated with Matrigel, in the presence of 5 ml of ‘neural induction medium’ containing DMEM/F12 supplemented with GlutaMax (1/1; Thermo-Fisher Scientific; Cat. No. 10565-018), Neurobasal medium (1/1; Thermo-Fisher Scientific; Cat. No. 21103-049), N2 (0.5X; Thermo-Fisher Scientific; Cat. No. 17504-044), B27 (0.5X; Thermo-Fisher Scientific; Cat. No.

17502-048), ascorbic acid (100 μM; Sigma-Aldrich; St. Louis, MO, USA; Cat. No. A5960), L-Glutamax (0.5X; Thermo-Fisher Scientific; Cat. No. 35050-061), antibiotic-antimycotic (1X; Thermo-Fisher Scientific; Cat. No. 15240-062), 3 μM CHIR99021 (Selleck Chemicals; Houston, TX, USA; Cat. No. S2924), 2 μM DMH1 (Selleck Chemicals; Cat. No. S7146), and 2 μM SB431542 (Selleck Chemicals; Cat. No. S1067), and 10 μM ROCK inhibitor (compound Y-27632 2HCl; Selleck Chemicals; Cat. No. S1049). After 24 hours, the medium was replaced with the same medium without ROCK inhibitor. The culture medium was changed every other day until DIV6, when induced NPCs were instructed to acquire a caudalized and ventralized progenitor cell identity as described.^33, 34^ Briefly, NPCs were dissociated with Gentle Cell Dissociation Reagent and split 1:6 in NPC expansion medium composed of the same medium described above, supplemented with retinoic acid (RA) (0.1 μM; Sigma-Aldrich; Cat. No. R2625) and purmorphamine (0.5 μM; Sigma-Aldrich; Cat. No. SML-0868) in combination with 1 μM CHIR99021, 2 μM DMH1 and 2 μM SB431542 reagents. The culture medium was changed every other day until DIV12, when they were split again 1:6 and expanded with the same medium containing 3 μM CHIR99021, 2 μM DMH1, 2 μM SB431542, 0.1 μM RA, 0.5 μM purmorphamine, and 500 μM valproic acid (VPA; Sigma-Aldrich; Cat. No. P4543) till DIV19. The ensuing caudalized and ventralized NPCs were validated by real-time polymerase chain reaction (RT-PCR) and immunocytochemistry.

### Differentiation of astrocytes from human iPSC-derived neural progenitor cells

Induced caudalized/ventralized NPCs were differentiated into astrocytes starting at DIV19 using a defined medium, essentially as described previously.^17^ NPCs were seeded at low cell density (15,000 cells/cm^2^) in two T25 flasks in the presence of 5 ml of NPC expansion medium containing ROCK inhibitor. Next day, medium was replaced with ‘Astrocyte Differentiation Medium 1’ [ScienceCell Astrocyte Growth Medium (ScienCell Research Laboratories; Carlsbad, CA, USA; Cat. No. 1801b) containing astrocyte growth supplement (ScienCell Research Laboratories; Cat. No. 1852), 1% fetal bovine serum (FBS) (ScienCell Research Laboratories; Cat. No. 0010), 50 U/ml penicillin G, 50 mg/ml streptomycin]. Cells were split 1:4 every week and were maintained under these culture conditions for 30 days. Half medium was replaced with fresh medium every 3 to 4 days. At DIV50, cultures were switched to ‘Astrocyte Differentiation Medium 2’ (same as Astrocyte Differentiation Medium 1 but lacking FBS), and routinely analyzed at DIV80. Induced astrocytes were validated by immunocytochemistry, RT-PCR, and calcium imaging.

### Characterization of induced cells by immunocytochemistry

Induced human NPCs and astrocytes were analyzed by immunocytochemistry, which was performed as described previously.^35^ The following primary antibodies were used: anti HOXA5 and anti-HOXC9 (1/67000, kindly provided by Dr. Jeremy Dasen, New York, University School of Medicine), rabbit anti-FOXG1 (1/300; Abcam; Cat. No. Ab196868), mouse anti-NKX6.1 (1/500; DSHB; Cat. No. F55A10), rabbit anti-PAX6 (1/500; BioLegend; Cat. No. 901301) mouse anti-GFAP (1/1,000; Sigma-Aldrich; Cat. No. G3893); mouse anti-S100B (1/500; Sigma-Aldrich; Cat. No. S2532); mouse anti-GAP JUNCTION PROTEIN ALPHA 1 (GJA1) (1/500; Abcam; Cat.

No. 11369), and rabbit anti-SLC1A2/EAAT2/GLT-1 (SLC1A2) (1/500; Abcam; Cat. No. 41621). Secondary antibodies against primary reagents raised in various species were conjugated to Alexa Fluor 555 and Alexa Fluor 488 (1/1,000; Invitrogen; Burlington, ON, Canada). Actin polymerization was visualized by staining of F-actin using Alexa-Fluor-488 phalloidin (1/500; Thermo Fisher; Cat. No. A12379). Images were acquired with a Zeiss Axio Observer Z1 Inverted Microscope using 20X magnification (N.A 0.8) and a ZEISS Axiocam 506 mono camera.

### Characterization of induced cells by real-time polymerase chain reaction

RNA extraction and real-time polymerase chain reaction (RT-PCR) were performed as described.^36^ Analysis of gene expression was conducted using the following oligonucleotide primers: Taqman probes *FOXG1*, Hs01850784_s1; *HOXA3*, Hs00601076_m1; *HOXA5*, Hs00430330_m1; *HOXB8*, Hs00256885_m1; *NKX2.2*, Hs00159616_m1; *NKX6.1*, Hs00232355_m1; *NKX6.2*, Hs00752986_s1; *PAX6*, Hs01088114_m1; *PAX3*, Hs00240950_m1; *PAX7*, Hs00242962_m1; *GFAP, Hs00909233_m1*; *S100B*, Hs00902901_m1; *CD44*, Hs01075864_m1; *GJA1*, Hs00748445_s1; *SLC1A2*, Hs01102423_m1; *SLC1A3/EAAT1/GLAST (EAAT1), Hs00904823_g1; AQUAPORIN-4 (AQP4), Hs00242342_m1; TFAP2A*, Hs01029413_m1; *REELIN*, Hs01022646_m1; *POTASSIUM INWARDLY RECTIFYING CHANNEL SUBFAMILY J MEMBER 10* (*KCNJ10*), Hs00158426_m1; *AMIGO2*, Hs00827141_g1; *SERGLYCIN*, *Hs01004159_m1; COMPLEMENT C1S (C1S*), Hs00156159_m1; *COMPLEMENT C3* (*C3*), Hs00163811_m1; *INTERLEUKIN-1BETA* (*IL-1β), Hs01555410_m1; TUMOR NECROSIS FACTOR ALPHA* (*TNF-α), Hs 00174128_m1; C-C MOTIF CHEMOKINE LIGAND 2* (*CCL2*), Hs00234140_m1; C-X-C MOTIF CHEMOKINE LIGAND 10 (*CXCL10*),

Hs00171042_m1). Primer/probe sets were obtained from ThermoFisher Scientific. Data were normalized with *BETA-ACTIN* and *GAPDH* (*ACTB Hs01060665_g1; GAPDH Hs02786624_g1*). Relative quantification (RQ) was estimated according to the ΔCt method.^37^

### Detection of calcium transients in human iPSC-derived astrocytes

DIV80 human iPSC-derived astrocytes were incubated with the calcium indicator Fluo-4 AM (1μM; ThermoFisher Scientific; Cat. No. F14201) for 30 minutes, followed by two rinsing steps with ScienCell Astrocyte Growth Medium, supplemented with antibiotics, for 10 min each. Time-lapse microscopy was then performed at 0.5 Hz with a Zeiss Axio Observer Z1 Inverted Microscope using 20X magnification (N.A 0.8) and a ZEISS Axiocam 506 mono camera.

Temperature (37°C), humidity and CO_2_ (5%) were maintained constant throughout the course of the experiments through TempModule S and CO_2_ moduleS devices (PeCon GmbH; Erbach, Germany). LED 488 nm was set at 20% and exposure time was 200 ms to avoid phototoxicity. For spontaneous calcium wave acquisition, recording was conducted for 5 min and focus stabilisation was achieved with the use of Zeiss Definite focus every 30 images. For induced calcium wave acquisition, recording was performed for 1 min, before the addition of 10 μM ATP, followed by an additional 4 min recording. Analysis of at least 25 cells per acquisition video was performed using the calcium signaling analyzer tool, CaSiAn.^38^

## Results

### Generation of caudalized and ventralized neural progenitor cells from human iPSCs

To generate human iPSC-derived NPCs with the potential to give rise to cells with properties of ventral spinal cord astrocytes, iPSCs were first instructed to undergo neural induction *in vitro* by inhibiting BMP/TGFβ signaling pathways for 6 days using a combination of the small molecules CHIR99021, DMH1 and SB431542. The ensuing NPCs were exposed to ventralizing and caudalizing cues by combined treatment with 0.5 μM purmorphamine and 0.1 μM retinoic acid (RA) for 6 additional days, as previously described^33, 34^, followed by NPC expansion till DIV19 (Fig. 1A).

**Figure 1.**
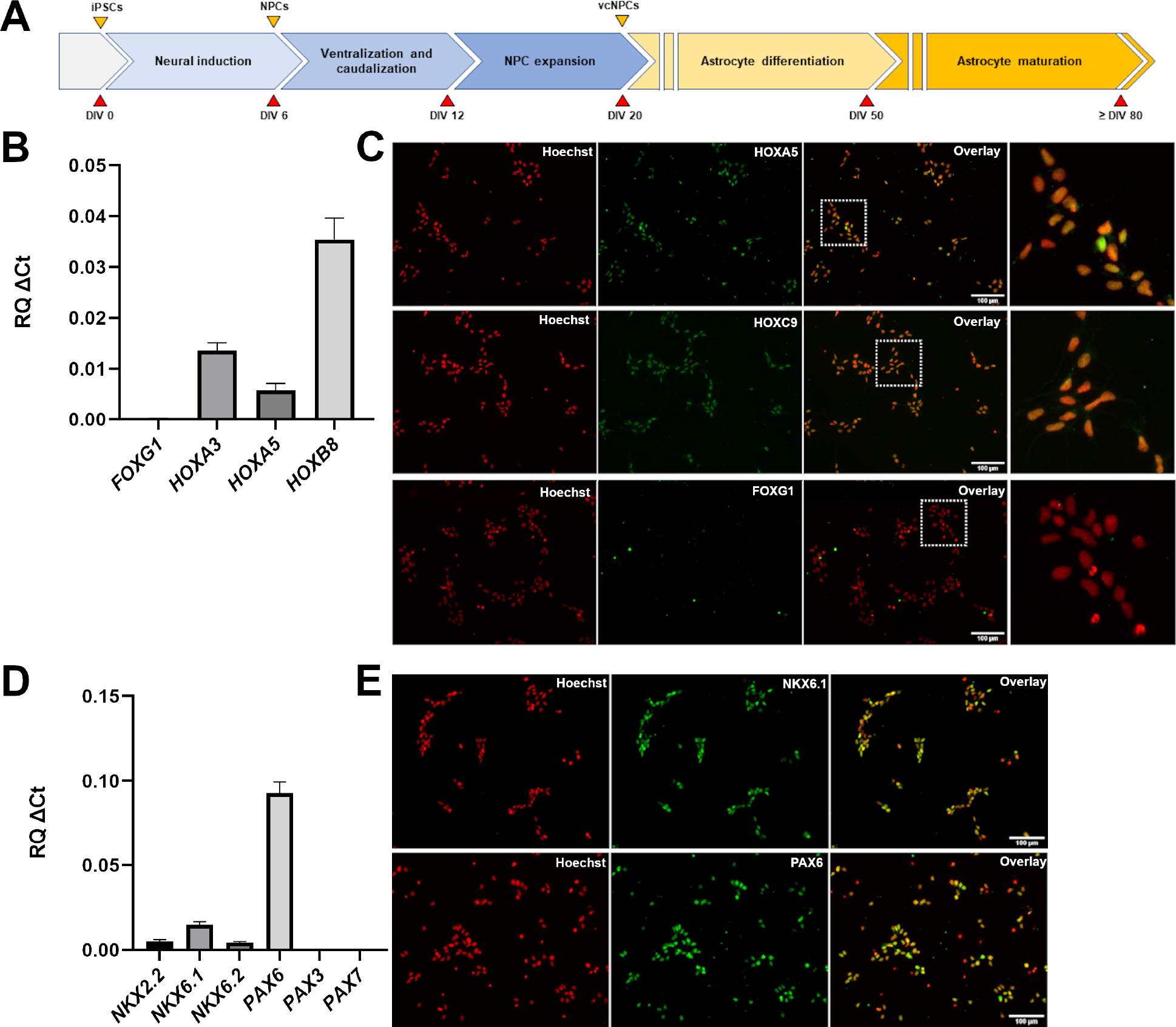
Generation and characterization of ventralized and caudalized neural progenitor cells. **A**) Schematic representation of the main steps of the differentiation protocol used to generate caudalized and ventralized neural progenitor cells and astrocytes. Days *in vitro* (DIV) are indicated. NPCs, neural progenitor cells; vcNPCs, ventralized and caudalized neural progenitor cells. **B**) Real-time PCR analysis of the expression of forebrain marker *FOXG1* and caudal markers *HOXA3*, *HOXA5*, and *HOXB8* in NPCs at DIV19. **C**) Representative images of immunofluorescence analysis of induced NPCs, showing that most of the cells are positive for HOXA5 and HOXC9 (green), but negative for FOXG1, at DIV19; Hoechst counterstaining (red) is shown. In each row, dotted-line box in the third panel indicates area shown at higher magnification in the fourth panel on the right-hand side. **D**) Real-time PCR analysis of the expression of a panel of ventral-to-dorsal spinal cord markers, namely *NKX2.2*, *NKX6.1*, *NKX6.2*, *PAX6*, *PAX3* and *PAX7*. **E**) Representative images of immunofluorescence analysis of induced NPCs, showing that most cells are positive for NKX6.1 and PAX6 (green) at DIV19; Hoechst counterstaining (red) is shown.

Immunocytochemistry and RT-PCR studies were performed to confirm the generation of NPCs with caudal (eg, brainstem/spinal) and ventral neural tube properties. RT-PCR showed that three transcripts associated with caudal positional identity, such as *HOXA3*, *HOXA5*, and *HOXB8*^39–41^, were expressed in NPCs exposed to purmorphamine and RA (‘purmorphamine/RA’ hereafter) (Fig. 1B). To complement this observation, we examined the expression of *FOXG1*, a marker of forebrain NPCs.^42^ *FOXG1* transcript level was below the limit of detection (Fig. 1B). In agreement with these findings, immunocytochemistry showed that the majority of NPCs expressed the proteins HOXA5 (96.1% ±1.5%) and HOXC9 (another caudal *HOX* gene product) (99.3% ± 1.2%), but were negative for nuclear FOXG1 expression (Fig. 1C). To validate the ventral identity of NPCs exposed to purmorphamine/RA, we examined the expression of genes marking ventral and dorsal NPCs in the spinal cord *in vivo*, including *NKX2.2*, *NKX6.1*, *NKX6.2*, *PAX6*, *PAX3* and *PAX7*.^43–45^ This analysis showed detectable expression of the exclusively, or predominantly, ventral markers *NKX2.2*, *NKX6.1*, *NKX6.2*, as well as *PAX6*, which marks both ventral and dorsal spinal cord progenitor domains *in vivo*; in contrast, the predominantly dorsal markers *PAX3* and *PAX7* were undetectable (Fig. 1D). Immunocytochemistry confirmed that most of the induced NPCs expressed NKX6.1 (91% ±1.5%) and PAX6 (93.3% ± 1.9%) (Fig. 1E). Together, these results demonstrate robust generation of NPCs with ventral and caudal characteristics from human iPSCs by DIV19.

### Generation of astrocytes from caudalized and ventralized neural progenitor cells

NPCs characterized as described above were induced to undergo astrocytic commitment and differentiation *in vitro* starting at DIV20 (Fig. 1A). Expression of common astrocyte marker genes, such as *GFAP*, *S100B*, and *VIMENTIN*, was detected in differentiating cultures as early as DIV50 (not shown). At DIV80, most induced cells displayed a fibrous morphology characterized by long and thin extensions (Fig. 2A) and were positive for S100B expression (Fig. 2B). Numerous S100B- positive cells exhibited robust co-expression of GFAP, with fewer cells displaying lower or undetectable GFAP levels (Fig. 2B). Induced cells expressed other typical astrocyte marker genes, such as *GJA1* (also referred to as *CONNEXIN-43*), *SLC1A2* (also known as *EXCITATORY AMINO ACID TRANSPORTER 2* - *EAAT2*), *SLC1A3* (a.k.a., *EAAT1*), and *CD44* (Fig. 2C-E), confirming their astrocytic molecular identity.

**Figure 2.**
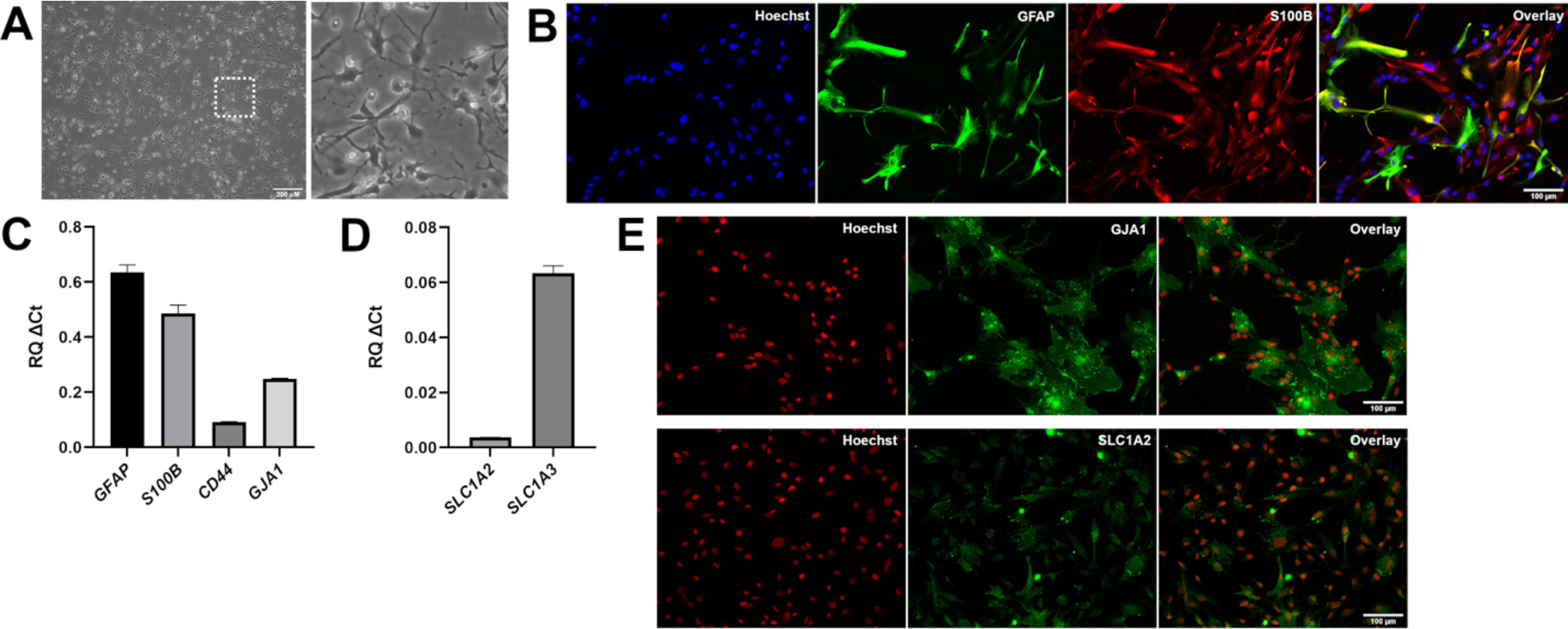
Characterization of astrocytes generated from ventralized and caudalized neural progenitor cells. **A**) Phase contrast image of DIV80 astrocytes derived from ventralized and caudalized NPCs, showing that most cells have stellate-like morphologies with long and thin extensions. Box in left-hand panel indicates area shown at higher magnification in right-hand panel. **B**) Representative double-labeling immunofluorescence analysis of GFAP (green) and S100Β (red) expression in DIV80 astrocytes; Hoechst counterstaining (blue) is shown. The majority of induced cells express S100B, and a significant proportion of S100B-positive cells co- express GFAP at high level, while other S100B-positive cells co-express GFAP at low level. **C, D**) Real-time PCR analysis of the expression of *GFAP*, *S100B*, *CD44*, *GJA1* (**C**), as well as *SLC1A2* and *SLC1A3* (**D**) in induced astrocytes. **E**) Immunofluorescence analysis of GJA1 and SLC1A2 (green) expression in DIV80 astrocytes generated from ventralized and caudalized NPCs; Hoechst counterstaining (red) is shown.

The caudalized nature of the astrocytes generated from purmorphamine/RA-treated NPCs was demonstrated by the persistent expression of *HOXA3*, *HOXA5*, and *HOXB8*, and lack of rostral marker *FOXG1* expression (Fig. 3A). The continued expression in spinal astrocytes of *HOX* genes first expressed in their progenitors is consistent with results of previous *in vitro* studies.^41^

**Figure 3.**
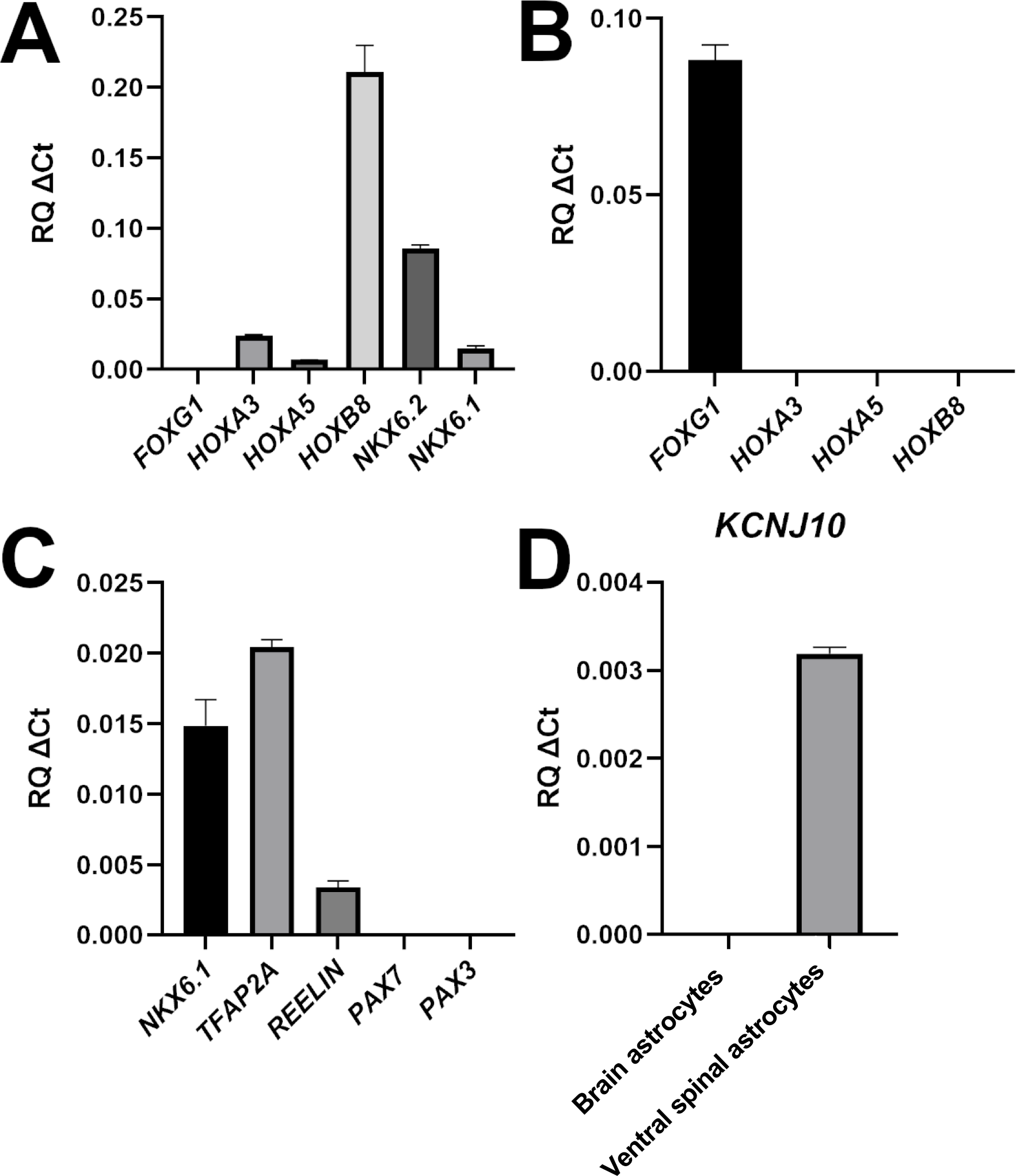
Characterization of human iPSC-derived VS astrocytes. **A**, **B**) Real-time PCR analysis of the expression of the indicated markers of rostrocaudal identity in either ventral spinal (**A**) or brain (**B**) astrocytes. (**C**) Real-time PCR analysis of the expression of markers of ventral spinal identity in ventral spinal astrocytes. **D**) Comparison of *KCNJ10* expression in brain or ventral spinal astrocytes.

Conversely, forebrain astrocytes generated as described in^36^ expressed *FOXG1* but not *HOXA3*, *HOXA5*, and *HOXB8* (Fig. 3B). In agreement with these observations, astrocytes generated from purmorphamine/RA-treated NPCs also continued to express *NKX6.2* (Fig. 3A), which has been previously identified as a marker of spinal cord, but not forebrain, astrocytes.^41^ To validate the ventral phenotype of the induced astrocytes, we examined the genes *NKX6.1*, *REELIN*, and *TFAP2,* whose expression was shown to be preferentially associated with ventral, but not dorsal, spinal cord astrocytes.^41, 46^ All of these genes were expressed in astrocytes generated from purmorphamine/RA-treated NPCs; in contrast, no detectable expression of the dorsal markers *PAX3* and *PAX7* was observed (Fig. 3C). To extend this analysis, we examined the expression of the gene encoding KCNJ10, the human counterpart of the murine inward-rectifying potassium channel Kir4.1, which is preferentially expressed in astrocytes located in the ventral horn of the rodent spinal cord *in vivo*.^47, 48^ Astrocytes generated from purmorphamine/RA-treated NPCs were positive for *KCNJ10* expression; in contrast, brain astrocytes generated in the absence of ventralizing cues^36^ displayed low or undetectable *KCNJ10* expression (Fig. 3D). Together, these results provide evidence for robust generation from human iPSCs of astrocytes with molecular features resembling those of ventral spinal cord astrocytes. These induced cells will be referred to hereafter as human iPSC-derived ‘ventral spinal-like’ (VS) astrocytes.

### Lack of overt signs of activation in human iPSC-derived VS astrocytes

Human iPSC-derived astrocyte preparations with the best potential to offer insight into mechanisms of astrogliosis should not be intrinsically reactive as a result of *in vitro* derivation conditions. The observation that human iPSC-derived VS astrocytes express *KCNJ10*, which was shown to be down-regulated in reactive astrocytes in multiple studies^49–51^, suggested that these astrocytes were not reactive. To examine this possibility further, we determined the expression levels of a variety of genes previously shown to be highly expressed in reactive astrocytes. During astrogliosis, there is a negative correlation between the expression of KCNJ10 (Kir4.1 in rodents) and AQUAPORIN-4 (AQP4), an astrocytic water channel. Both KCNJ10 and AQP4are mainly localized to astrocytic endfeet and play important roles in potassium homeostasis and blood-brain barrier integrity.^52, 53^ In contrast to the down-regulation of KCNJ10 during astrogliosis^49–51^, AQP4 levels increase in reactive astrocytes.^52–54^ Importantly, such an inverse correlation was also observed during ALS-associated astrogliosis.^56^ RT-PCR analysis showed that human iPSC-derived VS astrocytes expressed very low levels of *AQP4* (Fig. 4A), an observation consistent with the results of previous *in vitro* analysis of iPSC-derived spinal astrocytes.^41^ This finding also suggests that human iPSC-derived VS astrocytes are not intrinsically activated.

**Figure 4.**
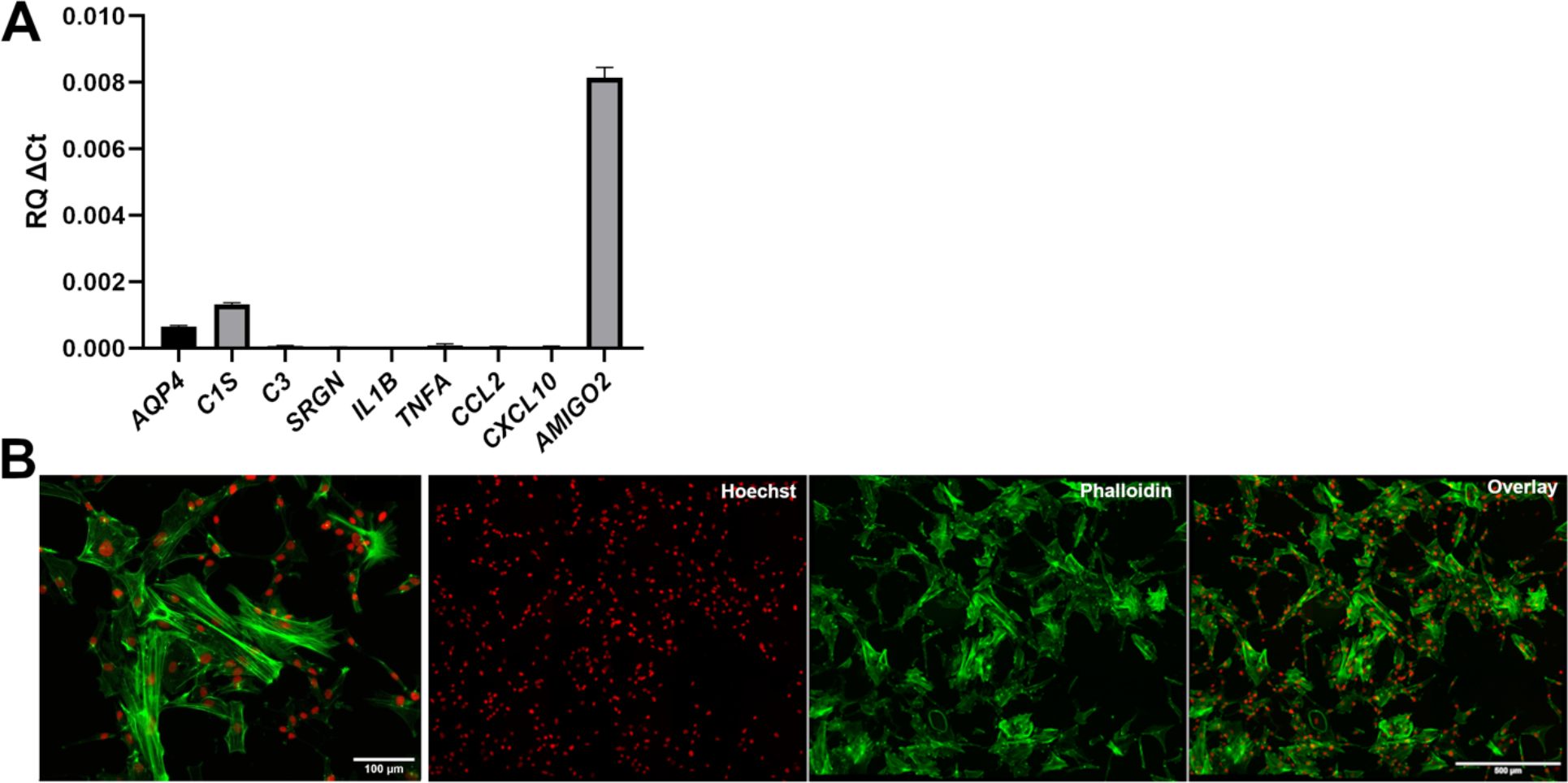
Lack of detectable signs of activation in human iPSC-derived ventral spinal astrocytes. **A**) Real-time PCR analysis of the expression of a panel of astrogliosis markers. **B**) Representative images, at different magnifications, of the actin cytoskeleton of induced astrocytes visualized by Alexa488-conjugated phalloidin staining (green); Hoechst counterstaining (red) is shown.

To extend this analysis, we next examined the expression of a number of genes associated with a reactive astrocyte phenotype *in vivo* and *in vitro*. These included complement pathway components such as *C1S* and *C3*, *SRGN*, *IL1B, TNFA, CCL2*, *CXCL10* and *AMIGO2*, all of which were reported to be up-regulated in reactive astrocytes.^57–61^ RT-PCR analysis showed little or no detectable expression of most of these genes in human iPSC-derived VS astrocytes, with the exception of *AMIGO2* (Fig. 4A). The latter observation is consistent with the physiological expression of *AMIGO2* in spinal cord astrocytes.^62^ Of note, the levels a number of pro-inflammatory cytokines were up-regulated when human iPSC-derived VS astrocytes were exposed to a pro-reactive astrocyte phenotype medium containing TNF-α, IL-1α, and C1q^58, 59^, suggesting that these cells have the potential to become activated under the appropriate conditions (data not shown).

Based on these findings, we next investigated the cytoskeletal organization of human iPSC-derived VS astrocytes. It is known that F-actin stress fibers disassemble into a more disorganized G-actin network during astrogliosis, resulting in altered morphologies, also characterized by the presence of ring-like structures, ruffles, and radial actin filaments extending towards the astrocyte periphery.^63, 64^ DIV80 human iPSC-derived VS astrocytes displayed an organized network with discernable F-actin stress fibers using phalloidin staining (Fig. 4B). Taken together, these results provide evidence suggesting that human iPSC-derived VS astrocytes are not activated under the examined experimental conditions.

### Human iPSC-derived VS astrocytes display spontaneous and ATP-induced calcium waves

To further characterize the properties of human iPSC-derived VS astrocytes, and to assess their potential for the study of mechanisms of astrocyte-neuron communication, we next examined whether these preparations would exhibit spontaneous and induced cytosolic calcium [Ca^2+^] concentration transients. Calcium concentration waves are a physiological property of functional astrocytes and represent one of the mechanisms by which astrocytes regulate neuronal functions, through the release of ‘gliotransmitters’.^65, 66^ We incubated DIV80 human iPSC-derived VS astrocytes with the Ca^2+^ indicator Fluo-4 AM, in the absence (spontaneous Ca^2+^ transients) or presence of 10 μM ATP (induced Ca^2+^ transients) (Fig. 5). Spontaneous Ca^2+^ waves were readily observed in the majority of cells, with signals frequently propagating from one cell to the adjacent ones (Fig. 5A, B; Suppl. Video 1 in Supplementary Information). Treatment with 10 μM ATP triggered a sharp increase in Ca^2+^ concentration, followed by multiple waves of calcium release, suggestive of Ca^2+^-induced Ca^2+^release (Fig. 5D, E; Suppl. Video 2 in Supplementary Information). We quantified Ca^2+^ transient characteristics, including amplitude, spike width, and inter-spike intervals, using CaSiAn, an open software tool.^38^ The characteristics of the spontaneous and ATP-evoked Ca^2+^ waves were consistent with previous studies^17, 36, 38, 41^, suggesting further that human iPSC-derived VS astrocytes resemble physiological astrocytes (Fig. 5C, F). Together, these findings provide evidence that human iPSC-derived VS astrocytes display properties, such as spontaneous and evoked Ca^2+^ oscillations, that are consistent with *in vitro* developmental maturation and suitability for the study of neuron-astrocyte communication mechanisms.

**Figure 5.**
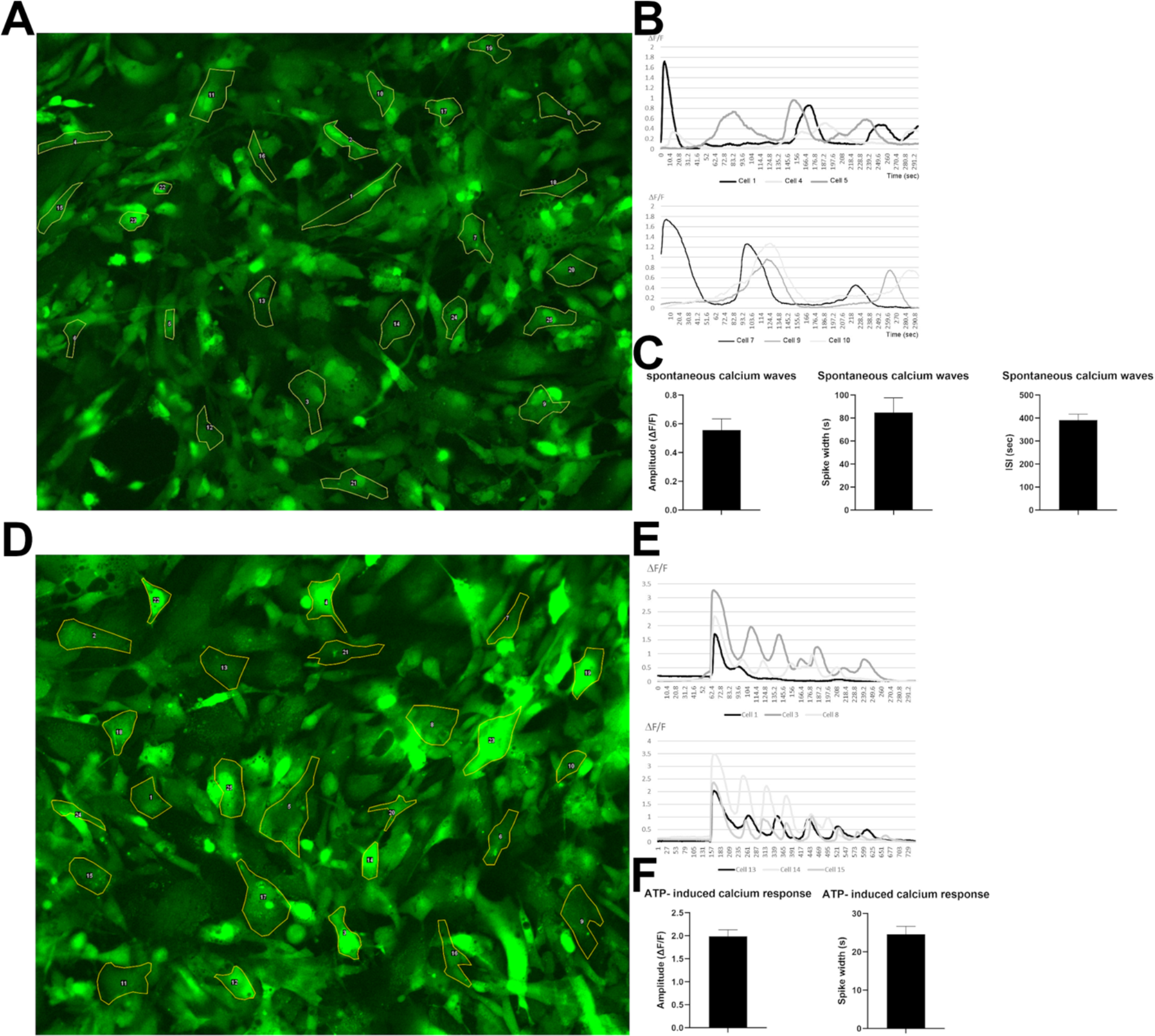
Spontaneous and induced calcium transients in human iPSC-derived VS astrocytes. **A**-**C**) Analysis of spontaneous Ca^2+^ waves. **A**) Representative Fluo-4 AM fluorescence. **B**) ΔF/F signals over time for 6 selected cells. **C**) Quantification of spontaneous transients’ characteristics including amplitude of signals, spike width, and interspike intervals. **D**-**F**) Analysis of ATP-induced induced Ca^2+^ waves. **D**) Representative Fluo-4 AM fluorescence. **E**) ΔF/F signals over time for 6 selected cells. **F**) Quantification of ATP-induced transients’ characteristics including amplitude of signals, spike width, and interspike intervals.

## Discussion

Human iPSC-derived neuronal and glial experimental model systems are becoming increasingly important to complement existing animal models to investigate the mechanisms leading to neuronal loss in neurodegenerative diseases, including ALS and other motor neuron diseases.^67–70^

The success of human iPSC-based studies depends in large part on the ability to generate the specific types of neuronal and glial cells that are most relevant to the disease under study. In this regard, investigations of astrocyte involvement in ALS pathophysiology require the *in vitro* derivation of specific types of astrocytes with molecular and physiological properties resembling those of the astrocytes that functionally interact with motor neurons impacted by this disease. For instance, astrocytes involved in biological cross-talk with upper or lower motor neurons are not functionally equivalent to astrocytes found in areas of the nervous system where ALS-impacted neurons are not present. This situation is particularly evident in the spinal cord, where only astrocytes found in the ventral horn, where motor neurons are located, have the potential to affect motor neuron function and survival, as opposed to dorsally located spinal astrocytes^31^. Thus, astrocyte differentiation protocols aimed at investigating mechanisms of non-cell autonomous motor neuron degeneration in ALS must be optimized to generate cultures enriched for the most disease-relevant astrocyte subtypes. In this work, we have described a robust and timesaving derivation from human iPSCs of astrocytes with properties resembling those of astrocytes found in the ventral spinal cord *in vivo* (‘human iPSC-derived VS’). These cells exhibit a number of properties that suggest that they will offer an enhanced experimental system for the study of spinal cord astrocyte biology and non-cell autonomous mechanisms of lower motor neuron degeneration in ALS.

### Human iPSC-derived ventral spinal-like astrocytes have properties of physiological astrocytes

The human iPSC-based derivation protocol described in this study yields induced cells expressing a complement of typical astrocytic markers as early as 50 days after the start of the derivation process. By DIV80, many induced cultures display expression of S100B, GFAP, SLC1A2, SLC1A3, as well as GJA1, the main astrocyte gap-junction protein. More importantly, these cultures are composed of astrocytes expressing a panel of genes found in more mature astrocytes of the ventral spinal cord, including *TFAP2A*, *REELIN*, and *KCNJ10*. Conversely, they lack detectable expression of more dorsal markers like *PAX3* and *PAX7*. Most induced astrocytes exhibit spontaneous and induced Ca^2+^ transients by DIV80, consistent with the properties of mature astrocytes capable of functional communication with neuronal cells.

The time required to obtain robust numbers of astrocytes with the above-summarized properties is relatively rapid when compared to previous studies describing the generation and characterization of ventral spinal cord astrocytes from human iPSCs. For instance, Bradley and colleagues have previously reported a ventral spinal astrocyte differentiation strategy from human iPSCs in which NPCs are first expanded in suspension for 5.5 months before differentiation to astrocytes in an adherent culture for one additional week, thus requiring almost 6 months to obtain astrocytes with the indicated properties.^41^ A somewhat faster protocol, generating spinal astrocytes in about 3 months, was described by Hall and coworkers^71^, but those studies, as well as others describing the generation of spinal astrocytes (e.g., references 27, 72, 73), did not provide a characterization of the specific ventral features of the induced astrocytes. The availability of a streamlined derivation protocol for generating characterized ventral spinal astrocytes from human iPSCs will therefore offer an enhanced opportunity to specifically investigate mechanisms of spinal cord astrocyte biology in health and disease.

### Human iPSC-derived ventral spinal-like astrocytes provide an improved experimental system for lower motor neuron disease research

A number of *a priori* requirements need to be fulfilled by human iPSC-derived astrocytes before they can be deemed suitable for the study of astrocyte involvement in lower motor neuron degeneration in ALS, as well as other lower motor neuron diseases. At a minimum, these cells must exhibit molecular hallmarks of ventral brainstem/spinal cord astrocytes, because the latter are the specific astrocyte subtypes that functionally interact with lower motor neurons. As already discussed, human iPSC-derived VS astrocytes fulfill this requirement.

Lower motor neuron disease-relevant astrocyte preparations should also display defined biological properties in order to qualify as an informative experimental system to investigate astrocyte-associated mechanisms of ALS pathophysiology. As a start, they should facilitate the investigation of genes whose altered expression has been associated with ALS phenotypes. For instance, the expression of two channels found on astrocytic endfeet, the inward rectifier-type potassium channel family member, KCNJ10A, and the water channel, AQP4, is affected in opposite ways in animal ALS models, where KCNJ10A levels are decreased whereas AQP4 levels are increased.^55, 56, 74–76^ Consistently, KCNJ10 is downregulated in astrocytes derived from ALS patients with mutations in the *SOD1* gene.^48^ The abnormal expression of AQP4 has been associated with altered blood-brain barrier integrity in ALS sufferers, as well as impaired potassium homeostasis and glutamate dysregulation.^76, 77^ Based on these observations, we examined the expression of both *KCNJ10* and *AQP4* in human iPSC-derived VS astrocytes. We observed that these preparations express low levels of *AQP4*, suggesting that they should be suited to investigate the functional impact of *AQP4* up-regulation, a situation that would be expected in astrocytes generated from iPSCs derived from ALS patients. Conversely, human iPSC-derived VS astrocytes express detectable levels of *KCNJ10*, implying that they should provide an informative experimental system to investigate the functional consequence of *KCNJ10* down-regulation, as observed in ALS astrocytes. Both of these lines of studies would be challenging if astrocytes expressing high levels of *AQP4* or low levels of *KCNJ10*, respectively, were used.

Another property that would be expected *a priori* for disease-relevant iPSC-derived astrocytes is the lack of intrinsic activation in the absence of internal or external insults. This feature would facilitate the investigation of abnormal astrogliosis mechanisms, such as those associated with ALS.^25–27^ The present work has suggested that human iPSC-derived VS astrocytes do not exhibit obvious signs of activation, as implied by the low or undetectable expression of a number of previously described astrogliosis markers, such as *SRGN*, *IL1B*, *TNFA*, *CCL2*, and *CXCL10*, as well as low levels of *AQP4*, as already described The presence of well-organized F-actin stress fibers is also consistent with a non-reactive phenotype.

In summary, the present findings provide evidence that human iPSC-derived VS astrocytes have a mature phenotype characterized by the expression of genes typical of differentiated ventral spinal astrocytes, display both spontaneous and induced Ca^2+^ signaling, without exhibiting obvious signs of activation. The availability of a protocol enabling relatively rapid generation from human iPSCs of astrocytes with properties resembling those of astrocytes in the ventral spinal cord will begin to address the challenges brought about by the demonstrated functional heterogeneity of astrocytes in the brain and spinal cord^4, 6, 8^, the selected contributions of different types of astrocytes to neuronal degeneration in ALS and other neurodegenerative disease^78–80^, and the need for fast and robust derivation strategies yielding disease-relevant preparations of specific astrocyte subtypes.

## Acknowledgements

We thank Louise Thiry, Nisha Pulimood, Yeman Tang, Rita Lo, Gilles Maussion, and Valerio Piscopo for discussions, advice and assistance.

## Authors’ contributions

VS performed all microscopy experiments, data analysis, and figure preparation. VS and LG performed qPCR experiments. MC, GH and AF induced iPSCs into neural progenitor cells. VS performed astrocyte differentiation and maturation. SS, TMD, VS, GR conceived overall study plan. SS, TMD supervised the study. SS and VS wrote the manuscript.

